# The need to harmonize insecticide resistance testing: methodology, intensity concentrations and molecular mechanisms evaluated in *Aedes aegypti* populations in Central America and Hispaniola

**DOI:** 10.1101/2020.02.25.964270

**Authors:** Sarah Ledoux, Carolina Torres Gutierrez, Neil F. Lobo, Elizabeth Melany Murillo, Silvia Pérez, Rocío Guerra, Sayra Chanquin Avendano, Ángel Gabriel Orellana Herrera, Aarón Mendoza, Denis Escobar, Gavino Guzmán Contreras, Magdiel Rivera, Gilda Ventura, Rodrigue Anagonou, Eliane Pierre-Louis, Carmen Yurrita, Francisco J. López Hun, Camilo Duque, Eduardo Romero, Diane D. Lovin, Joanne M. Cunningham, Dereje Dengela, Allison Belemvire, Kellie Stewart, Nelson Grisales

## Abstract

**Background:** The Zika AIRS Project, a USAID-funded initiative worked across the Latin America and Caribbean regions from 2016 to 2019, as an emergency to contain the spread of the Zika virus. All entomological records in the target countries showed wide distribution and high abundance of *Aedes aegypti* populations, however the susceptibility profiles of these insects to insecticides commonly employed by vector control campaigns were in most cases incomplete or inexistent. In close collaboration with the Ministries of Health of individual countries, Zika-AIRS teams conducted insecticide susceptibility testing of an array of insecticides in *A. aegypti* populations of each country. Procedures applied met the standard international protocols instructed by the World Health Organization and Centers for Disease Control and Prevention.

**Methodology and main findings:** The insecticides tested were selected under categories such as pyrethroids, organophosphates and carbamate. Results showed *A. aegypti* populations displaying high and widely distributed resistance to all pyrethroids across countries, tolerance to organophosphates and full susceptibility to a carbamate. Key inconsistencies between testing methods are presented and discussed. Additionally, four *kdr* mutations were analyzed to detect molecular mechanisms of insecticide resistance. The screening for *kdr* mutations suggested the widespread nature of V1016I mutation, linked to pyrethroid resistance in *A. aegypti* populations distributed and sampled in the above mentioned regions.

**Conclusions and perspectives:** This multi-country study contributes with updated information to the public health decision-makers across Central America and the Caribbean. This study provided training and established technical networks for more effective and sustainable insecticide surveillance programs. Most but not all records of insecticide resistance in *A. aegypti* were consistent between methodologies, thus inconsistent issues are discussed here to call for further improvement in procedures and convey more practical guidelines for surveillance teams in countries where *Aedes*-borne diseases are endemic.

**Author summary:** At the forefront of the fight against arboviruses transmission is the insecticide-based vector control. All countries in the Latin American and Caribbean region invest valuable resources from their limited budget to acquire and implement insecticide-based tools, with non-existent or weak insecticide resistance monitoring programs. Hence, the USAID-funded Zika AIRS Project (ZAP) collaborated with the Ministries of Health of multiple countries to update the profile of susceptibility to insecticides in *Aedes aegypti* populations. We found widespread resistance to pyrethroid and organophosphate insecticides, which account to almost 100% of the products available to control adult mosquitoes. As we used both of World Health Organization and Centers for Disease Control and Prevention standard methods, we found many similarities and some inconsistencies in the susceptibility profiles obtained for the very same vector populations. Additionally, we obtained insight on potential molecular mechanisms of resistance across the countries, finding the *kdr* mutation V1016I possibly involved in loss of susceptibility.

This study is the biggest cross-country update of insecticide resistance for *Aedes aegypti* in years, and it should be used as evidence for improving the selection of insecticides in these countries and a call for further support to maintain insecticide resistance monitoring programs.

## Introduction

Arboviruses are the most widely transmitted vector-borne diseases in the world. It is estimated that dengue fever, chikungunya, yellow fever and Zika infect more than 390 million humans per year (1, 2). At least 3.9 billion people in 128 countries are at risk of infection by dengue virus alone (3), and according to the World Health Organization (WHO) 3-4 million people were affected by Zika virus in the Americas during the 2016 outbreak (4). During the current year (2019), the Central American and Caribbean regions have faced periods of high dengue transmission, that have forced countries like Honduras and Jamaica declared public health warnings in their territories and displayed emergency responses to counter dengue outbreaks (PAHO records of dengue incidence include 369,609 cases from Central America and 21,115 cases from the Caribbean region, as reported in October 1^st^, 2019) (5). *Aedes aegypti*, the primary vector for all major arboviruses, is a container-breeding mosquito well adapted to domestic habitats located in the vicinity or inside human houses. Given the behavioral plasticity, rapid life cycle and invasive nature of *Ae. aegypti*, its distribution is virtually worldwide, in tropical and sub-tropical regions and in wide-ranging anthropic settings that include urban and rural areas. The expansion of *Ae. aegypti* will continue as climate change progresses, increasing the risk of arboviruses transmission in the near future, even in temperate regions (6, 7).

Efforts in mosquito control include community education, environmental modifications (i.e. larval site management) and use of chemical insecticides (8), ideally within an integrated vector management (IVM) strategy (9). The application of chemical insecticides to eliminate *Ae. aegypti* at multiple developmental stages by targeting larval sites and adult female habitats is recommended in an IVM plan. Larviciding, as part of environmental management, may be applied using compression spraying, powder, or dissolved solid formulations (9). Adulticides are applied using residual surface treatments or spatial applications, where the former is recommended only for emergency events and the latter has both adulticide and larviciding effects (9).

Of the four WHO approved insecticide classes available for outdoor mosquito control (pyrethroids, organophosphates, neonicotinoids and carbamates) via ultra-low volume (ULV) spraying, only pyrethroids and organophosphates are widely used (10). This widespread and continuous use of a small number of insecticides has resulted in the emergence of insecticide resistance in wild *Ae. aegypti* populations, across entire regions of the Americas and other continents. Such occurrences have been reported by entomological monitoring programs across the globe with increasing frequency (11). Although vector control through ULV insecticide application remains as the preferred tool in Latin America, more evidence documenting its effectiveness is still required.

Regular surveillance generates the baseline evidence required for examining both intervention potential as well as efficacy. Local evidence should guide countries in the rational use of insecticides, and at the same time improve timing of operations and decisions involving type of applications required. Examples of strategies for vector control insecticide applications are rotations, mixtures or mosaic spraying. Despite campaigns by the Global Vector Control Response and Worldwide Insecticide Resistance Network, many countries still lack capacity – both technical and financial - to optimally mobilize vector control intervention strategies (12, 13). A key component of entomological surveillance programs that utilize IVM in public health systems, remains insecticide susceptibility testing on local mosquito vector populations (14).

To determine *Ae. aegypti* susceptibility to insecticides the World Health Organization and the Centers for Disease Control and Prevention (CDC, Atlanta, USA) have provided standard procedures for laboratory bioassays (15, 16). These two methodologies evaluate mosquito tolerance to insecticide-specific diagnostic doses over time. Although both procedures are widely accepted as laboratorial surveillance to determine the susceptibility status of mosquito populations to insecticides used in public health, there are limited comparisons of both procedures towards establishing concordance of results (17).

A second step in the insecticide surveillance procedures recommended by WHO (46) include that any mosquito population found to be resistant to a given insecticide(s) should be further exposed to higher concentrations in order to assess the strength of the phenotypic resistance originally documented with discriminating concentrations (i.e. the intensity of resistance). Procedures that evaluate the effect of synergists on the resistant phenotypes are also included. Furthermore, other techniques may elucidate the biochemical and molecular mechanisms of insecticide resistance.

The molecular mechanisms of insecticide resistance can be grouped into four main categories: 1) enhanced metabolic resistance, 2) mutations in target sites, 3) cuticular resistance and 4) behavioral resistance. From these, the most documented mechanism is the knockdown resistance (*kdr*), which is a target site mechanism that confers resistance to pyrethroids and organochlorines (18–21). In the Americas, *kdr* mutations have been reported in *Ae. aegypti* populations from Ecuador, (22), United States (23, 24), Colombia (25, 26), México (27–29), Brazil (30–35), Lesser Antilles(36–40), French Guiana (37), Venezuela (29, 41), Cuba (29, 39), Panamá (42) and Puerto Rico (43). Similar to insecticide resistance monitoring, molecular resistance research of *Ae. aegypti* populations is particularly limited for Central American and Caribbean countries (10, 26).

Though pyrethroids and organophosphates have been utilized for extended periods of time, even decades, to control *Ae. aegypti* in regions of Latin America and the Caribbean (LAC), very few countries have conducted regular surveillance on local mosquito populations to assess insecticide susceptibility. Only Mexico and Colombia have a consistent insecticide resistance monitoring program, with only Mexico reporting a nationwide study on *Ae. aegypti* insecticide susceptibility status in recent years (44).

This is the first multi-country study on *Ae. aegypti* insecticide resistance to a wide selection of insecticides products being used for vector control operations in El Salvador, Guatemala, Honduras, Dominican Republic and Haiti in recent years, in addition to exploring the molecular mechanisms expressed in wild populations of each country.

## Materials and methods

### Study sites

The study sites or sentinel sites selected were originally part of the United States Agency for International Development (USAID) funded Zika AIRS Project (ZAP) (45), implemented from 2016 to 2019 in order to combat the 2016 regional Zika emergency and reinforce vector control and monitoring capacity. The study sites were chosen in collaboration with each country’s Ministry of Health, and based on Zika incidence in epidemiological reports. All locations per country are listed in Table 1 and displayed in Figure 1.

**Figure 1.**
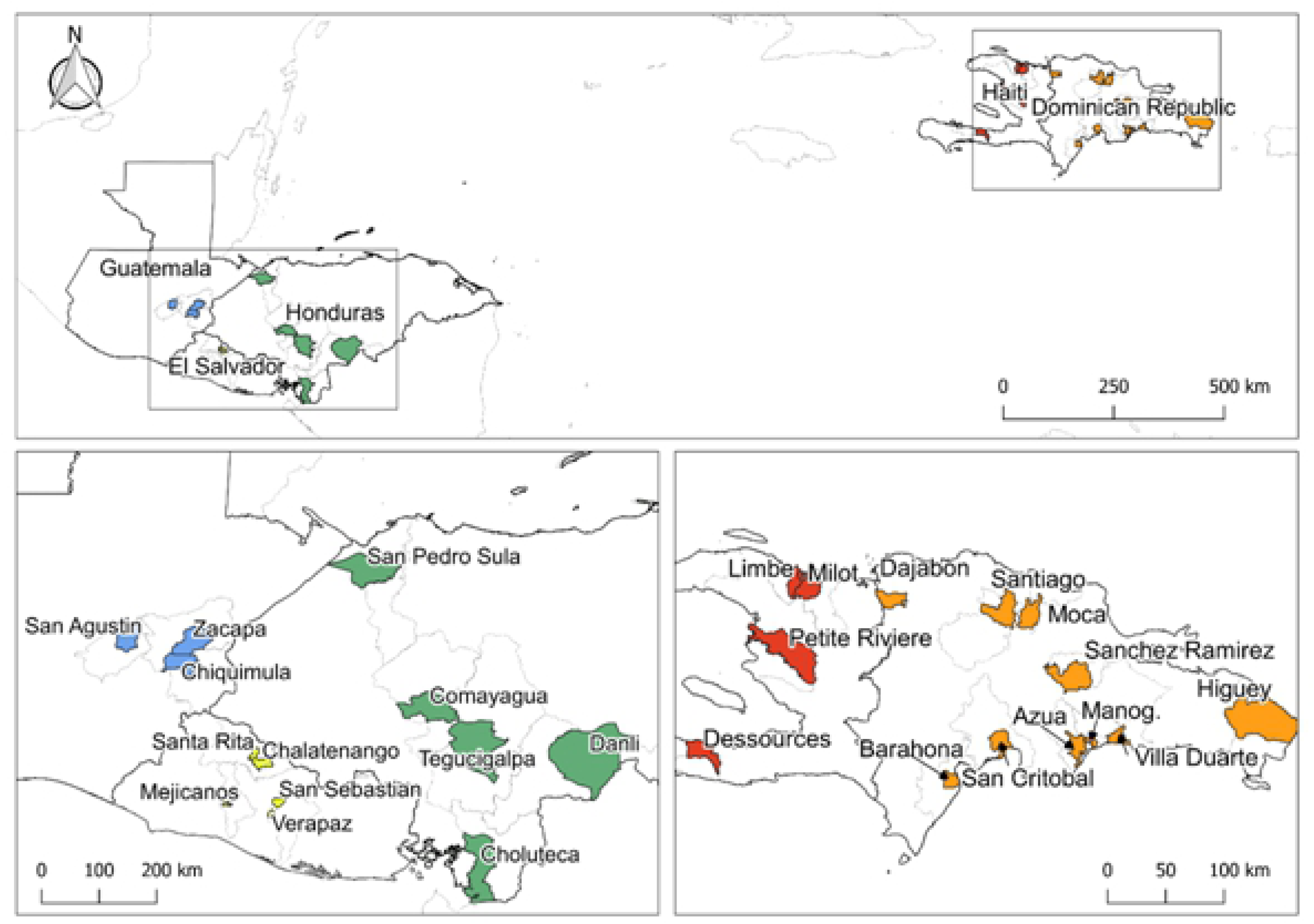
Countries and sentinel sites sampled for *Ae. aegypti* susceptibility tests to insecticides. The colored regions represent municipalities in Dominican Republic, El Salvador, Guatemala, and Honduras, and districts in Haiti. * The Dominican Republic site Manoguayabo was shortened to Manog.

**Table 1.**
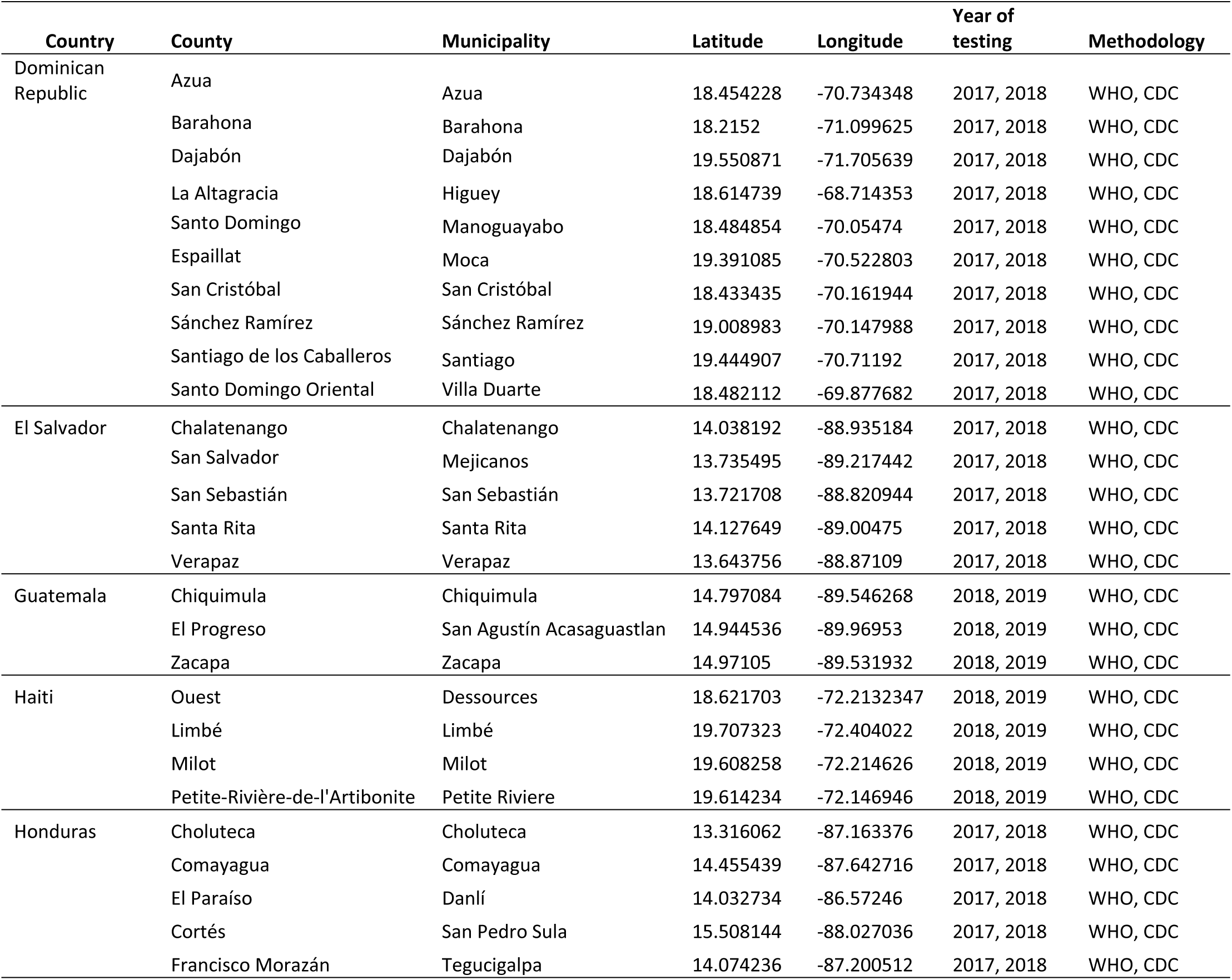
List of countries and study sites with geographical coordinates, year of testing and type of methodology conducted.

### Mosquito sampling

Mosquito collections and bioassays were performed throughout the duration of the Zika AIRS Project (2016 – 2019). Local populations of *Ae. aegypti* were sampled from ZAP’s sentinel sites in each country using two approaches: (i) larval collections from multiple houses and neighborhoods (n= 5-10), and (ii) ovitraps set in multiple premises per sentinel site (n= 5-10). Larval collections were performed by using pipettes and nets, and then transported to insectaries where specimens were reared to the adult stage. Larvae were fed with macerated fish or dog food pellets, with daily maintenance. Pupae were transferred to labeled mosquito cages representing each sentinel sites. Adults were fed *ad libitum* a 10% sucrose solution soaked in cotton balls. The insectary conditions recorded were 70% - 95% of relative humidity, and a temperature range from 26 °C – 29 °C and a photo-period of 12:12.

The ovitraps used were black plastic containers of approximately 1L of capacity, half-filled with 10% hay infusion and with the interior wall lined with a paper towel or germination paper as the oviposition substrate (adapted from (46)). Ovitraps were distributed in five to ten houses at least 200 meters apart. Once the oviposition papers were transferred to the laboratories, five to ten ovitrap papers with eggs were combined and immersed in dechlorinated water for hatching. Larvae, pupae and adult breeding conditions were identical to the ones described above. Adult mosquitoes were confirmed to be *Ae. aegypti* after the insecticide resistance tests using external morphological features described in taxonomical keys (47). F0 and F1 adult mosquitoes obtained under controlled insectary conditions were utilized for IR testing using WHO and CDC international standardized methodologies in each country with ZAP implemented entomological surveillance.

### Insecticides

The majority of the testing procedures were conducted between 2017 and 2018, with the exception of Haiti and Guatemala that completed the data during 2019. The detailed information on the exact months of collection and bioassays is available in the supplementary information (Supplementary information, Table S1).

All impregnated WHO papers with diagnostic concentrations (1x), intensity concentrations (5x and 10x the diagnostic dose), control papers and bioassay kits were obtained directly from the University Saints Malaysia (Penang, Malaysia). The standard insecticide-impregnated papers with the diagnostic doses used were permethrin (0.25 %), deltamethrin (0.03 %), lambda-cyhalothrin (0.03 %), etofenprox (0.5 %), alpha-cypermethrin (0.03 %), malathion (0.8 %), pirimiphos-methyl (0.21 %) and bendiocarb (0.1 %).

The insecticides and bioassay kits used for the CDC bottle assays were donated by the Centers of Disease Control and Prevention (Atlanta, United States) and included permethrin (15 µg/ml), deltamethrin (10 µg/ml), lambda-cyhalothrin (10 µg/ml), etofenprox (12.5 µg/ml), malathion (50 µg/ml), pirimiphos-methyl (20 µg/ml) and bendiocarb (12.5 µg/ml). A complete list of insecticides and concentrations is provided in the supplementary information (Table S2).

### WHO bioassays and CDC bottle assays

Three-to-five day old female mosquitoes were separated for at least one hour in paper cups before the bioassays. In WHO standard bioassays, the WHO protocol was followed using each insecticide’s diagnostic concentration for *Ae. aegypti* (48). At least four replicates with 25 mosquitoes each were used to test each insecticide, with at least one additional group exposed to control papers. After 60 minutes of exposure to insecticide, knockdown was recorded. Mortality was recorded 24 hours later.

The CDC standard IR bottle assays were performed according to the CDC guidelines (16). At least four replicates with 25 mosquitoes each were used to test each insecticide, with an additional group of 25 mosquitoes exposed only to the solvent in a separate bottle as a control. Knockdown was recorded every 15 minutes up to two hours, with the exception of the interval between 30 and 45 minutes where the readings were done every 5 minutes. The diagnostic time for all insecticides tested in this study was 30 minutes.

With the multi-country dataset per methodology, a qualitative comparison between methodologies and the status of mosquito populations tested (resistant vs. susceptible) was conducted. To summarize the level of alignment in the multi-country susceptibility records, the level of agreement between results obtained with WHO vs. CDC bioassays were classified as: “same” when exposed mosquitoes under both methodologies resulted either in i) resistance ii) suspected resistance or iii) susceptibility; “similar” when one assay resulted in resistance and the other test results in suspected resistance; and “different” when results were interpreted as resistant populations under one methodology and susceptible populations with the other, or suspected resistance, in one and susceptible in the other.

### WHO intensity bioassays

Standardized intensity bioassays (49) were adapted for *Aedes* with 5x and 10x the diagnostic concentrations of permethrin, deltamethrin and lambda-cyhalothrin, and were performed in El Salvador, Guatemala, Honduras and Haiti. Mortality values <98% with the 5x concentration indicates moderate resistance, while mortality values <98% with the 10x concentration suggest high intensity resistance.

### *Kdr* genotyping

A molecular screening for *kdr* mutations was conducted in order to characterize the allelic frequencies of four target site mutations incriminated in pyrethroid resistance of *Aedes aegypti* mosquitoes in Guatemala, El Salvador, Honduras, Haiti and Dominican Republic. Target mutations examined included I1011V, I1011M, F1534C and V1016I (11).

Molecular screening for *kdr* mutations I1011V, I1011M, F1534C and V1016I utilized DNA amplification and sequencing. For the DNA extraction the Rapid Alkaline DNA Extraction protocol was employed (50). Additionally, all amplification reactions included 25 µl total volume in 96-well PCR plates (Dot Scientific) in a Mastercycler Gradient thermocycler (Eppendorf). Each reaction contained 1X Taq buffer (50 mM KCl, 10 mM Tris pH 9.0, 0.1% Triton X), 1.5 mM MgCl_2,_ 200 µM dNTPs, 5 pmoles of each primer (except where noted), 1 unit of Taq DNA polymerase, and 3 µl of genomic DNA. PCR products were size fractionated by electrophoresis in 4% agarose gels stained with SybrSafe®, and visualized under UV light.

The mutation presence was characterized by using a primer multiplex to differentiate the wild-type from mutant individuals based on differences in amplicon size. The primers used for detecting each mutation are included as supplementary information (protocol for molecular testing in supplementary information, S3, S4). Sequencing of amplified fragments using both PCR primers was performed to confirm PCR results for the I1011V/M mutation (supplementary information, table S3.1).

For the molecular screening, the sample of *Ae. aegypti* individuals for each country consisted of 300 mosquitoes from Dominican Republic, and 150 individuals respectively for the rest of countries. For each lot of individuals, specimens were classified as survivors and non-survivors during the WHO tests using diagnostic concentrations of permethrin, lambda-cyhalothrin and deltamethrin. All molecular procedures were conducted in the Lobo Lab at the University of Notre Dame, Indiana, USA. The mosquito populations screened correspond toHiguey and Manoguayabo municipalities (Santo Domingo East) in Dominican Republic, Dessources in Haiti, San Sebastian in El Salvador, Tegucigalpa in Honduras and Zacapa city in Guatemala.

## Results

### WHO bioassays

*Aedes aegypti* populations showed widespread resistance to all five pyrethroids tested (alpha-cypermetrhin, deltamethrin, etofenprox, lambda-cyhalothrin, and permethrin) during the three-year monitoring program (Figure 2). Although mortality varied within and across countries, all populations demonstrated clear loss of pyrethroid susceptibility (<90% mortality). The intensity of the resistance to permethrin in El Salvador and Guatemala was high (10x the diagnostic concentration did not kill >98% of mosquitoes), while in Honduras resistance to permethrin was moderate (Figure 3). For lambda-cyhalothrin, Guatemala’s populations showed high intensity of resistance, while in Honduras the only population categorized as moderate resistance was San Pedro Sula. In El Salvador all the exposed populations displayed moderate resistance, except Santa Rita. Finally, deltamethrin resistance in the region ranged between low to high intensity (Figure 3).

**Figure 2.**
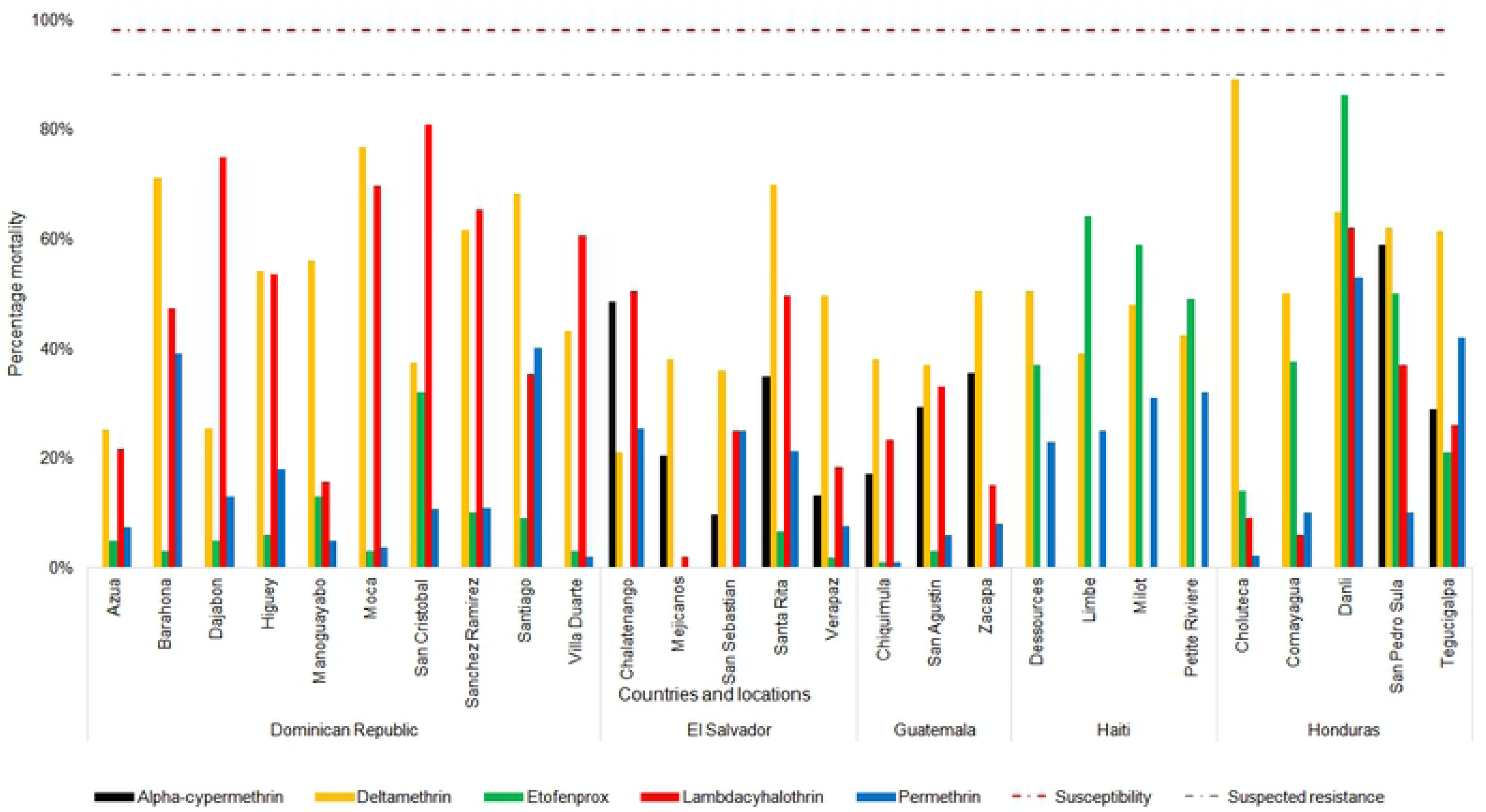
Mortality of *Ae. aegypti* to five pyrethroids obtained with WHO kits and diagnostic doses, the tested mosquito populations represent samples from Dominican Republic, El Salvador, Guatemala, Haiti and Honduras. The horizontal red dotted line represents 98% mortality which delimits the susceptibility threshold. Values between 98% and 90% mortality are interpreted as suspected resistance, and values below 90% mortality are interpreted as resistant to the corresponding insecticide. Note: alpha-cypermethrin was not tested in Dominican Republic and Haiti; lambda-cyhalothrin was not tested in Haiti.

**Figure 3.**
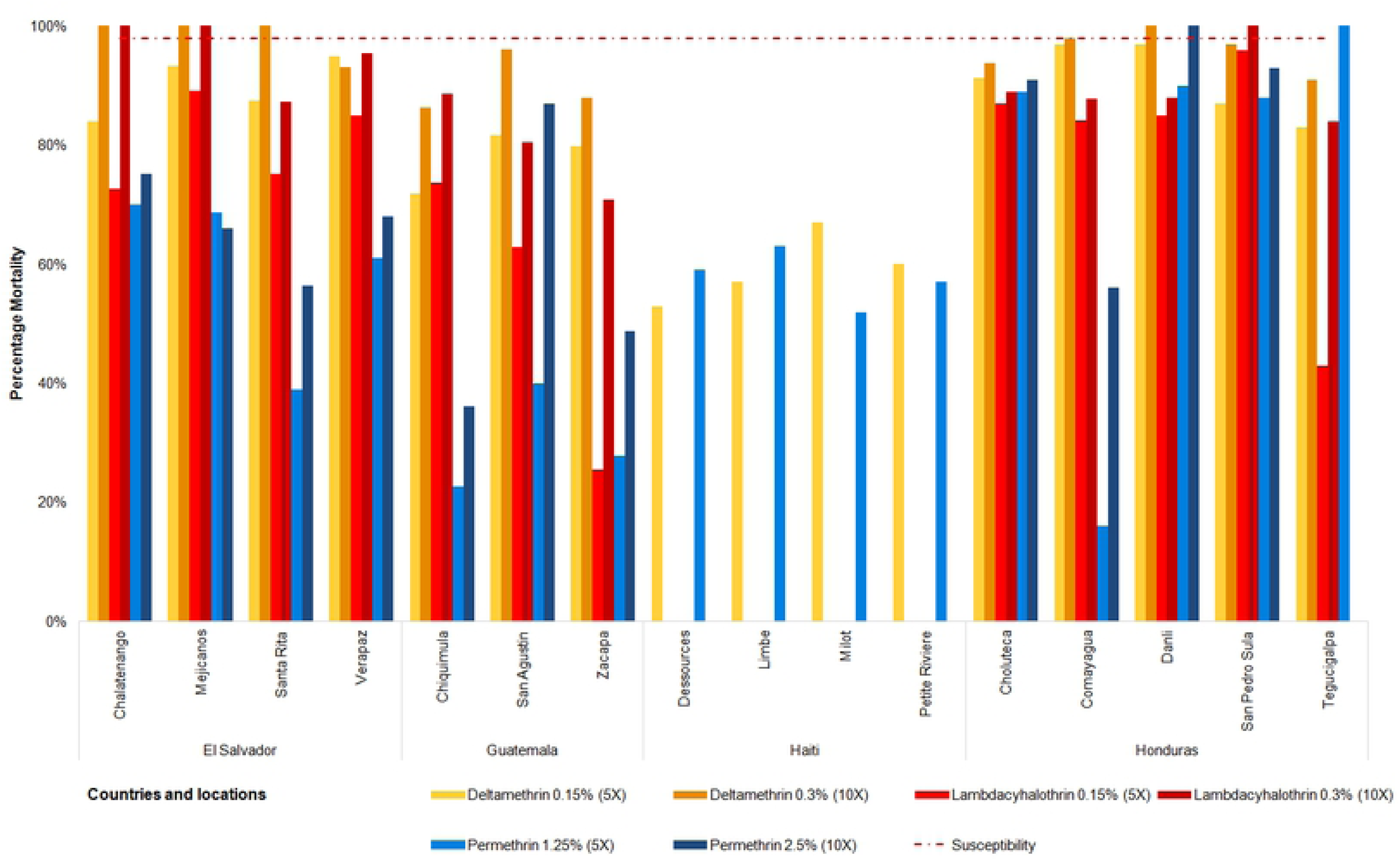
Intensity of resistance in *Ae. aegypti* populations from El Salvador, Guatemala, Haiti and Honduras to the WHO kits using three pyrethroids, with 5x insecticide concentrations (lighter color) and 10x concentrations (darker color). The horizontal red dotted line represents 98% mortality which delimits susceptibility. Note: 10x concentrations were not tested in *Ae. aegypti* populations from Haiti, nor in San Sebastian, El Salvador.

Resistance to the organophosphates malathion and pirimiphos-methyl was also documented. Only the *Ae. aegypti* population from one sentinel site in Guatemala, San Agustin, was susceptible to malathion. In the case of pirimiphos-methyl only mosquitoes from the West province in Haiti (Dessources and Petite Riviere), plus Higuey in Dominican Republic, resulted as susceptible (Figure 4).

**Figure 4.**
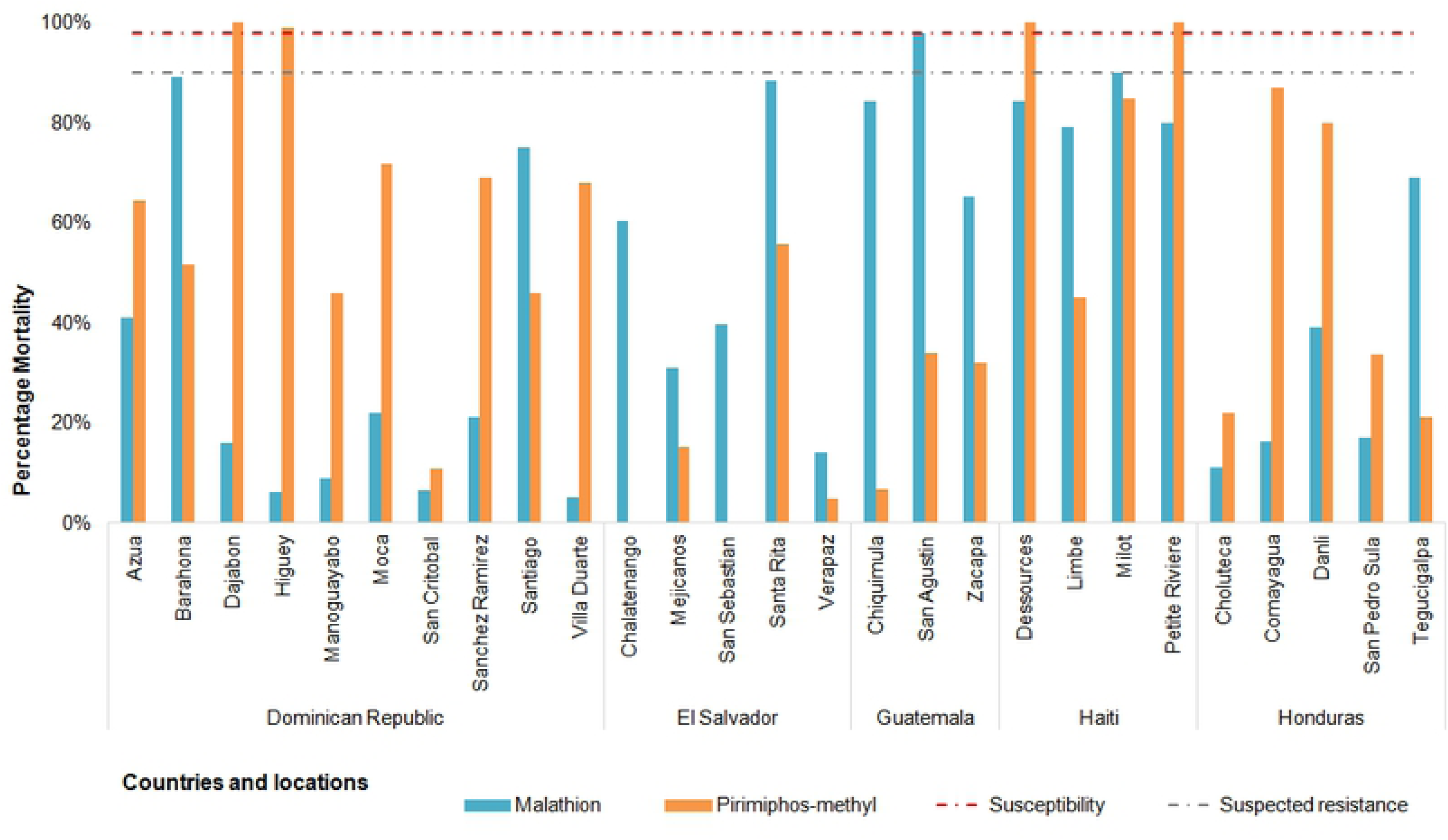
Mortality of *Ae. aegypti* populations from Dominican Republic, El Salvador, Guatemala, Haiti and Honduras when exposed to the organophosphates malathion and pirimiphos-methyl, using WHO methodology. The horizontal red dotted line represents 98% mortality and susceptibility threshold, between this threshold and the grey dotted line (90% mortality) values are interpreted as suspected resistance, and values below 90% mortality are interpreted as resistant to the corresponding insecticide.

Bendiocarb was only tested by WHO assays in the Dominican Republic, Guatemala and Honduras. In Honduras and Guatemala all *Ae. aegypti* populations tested demonstrated susceptibility or suspected resistance (further molecular tests need to confirm findings), except for the population from Chiquimula site in Guatemala. In contrast, all bioassays from Dominican Republic showed resistance or suspected resistance to bendiocarb (Figure 5).

**Figure 5.**
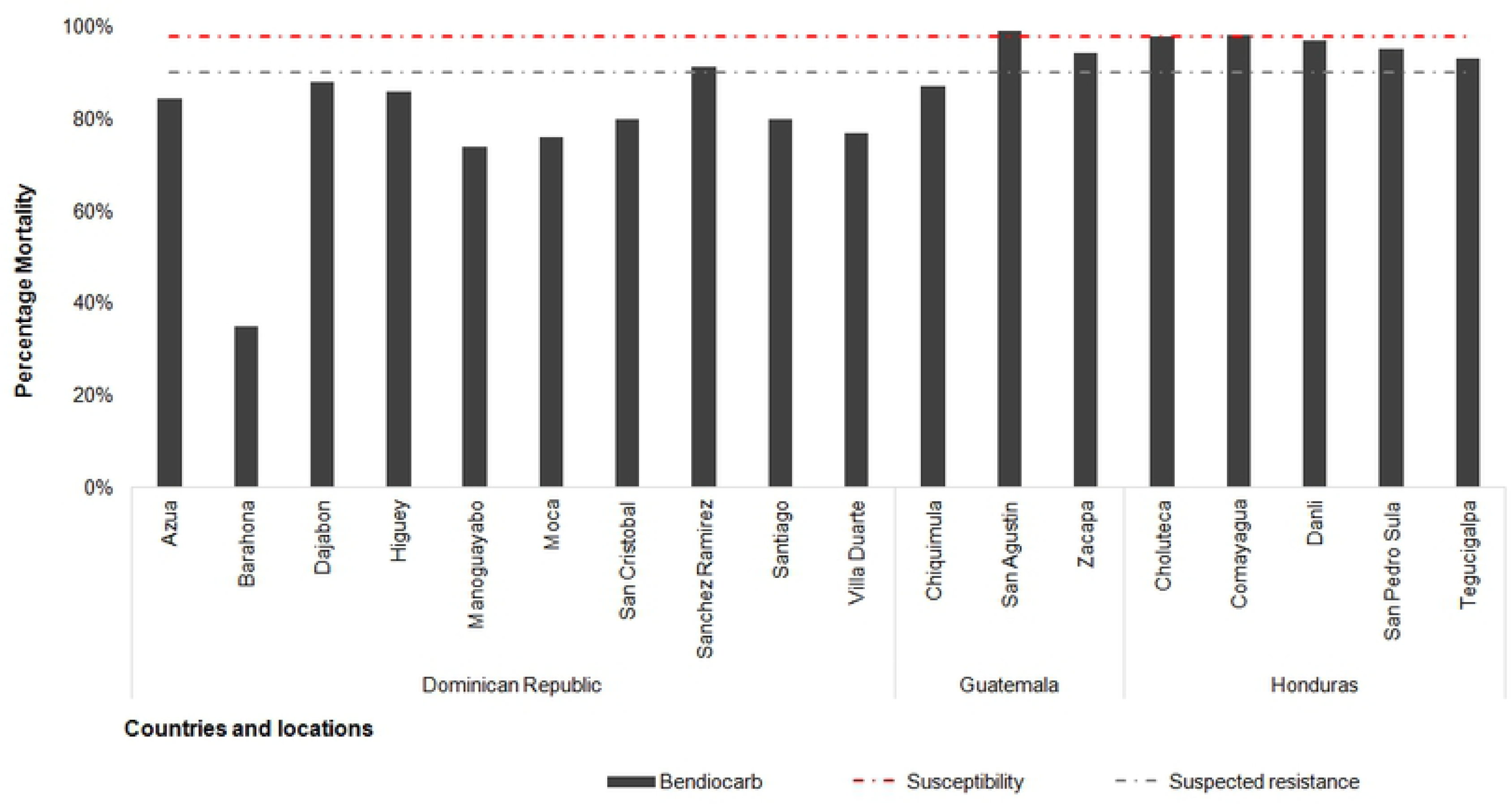
Mortality of *Ae. aegypti* populations from Dominican Republic, Guatemala, Haiti and Honduras when exposed to Bendiocarb, using WHO methodology. The horizontal red dotted line represents 98% mortality, and delimits the susceptibility threshold. Values between 98% mortality and the grey dotted line (90% mortality) are interpreted as suspected resistance, and values below 90% mortality are interpreted as resistant to the corresponding insecticide.

### CDC bottle bioassays

All mosquito populations from the Dominican Republic and Guatemala were susceptible to deltamethrin (Figure 6). While in Honduras, three of five sites showed resistance, in El Salvador all populations showed resistance (Figure 6). Lambda-cyhalothrin susceptibility was present in some populations of Dominican Republic and Honduras, but the majority of the *Ae. aegypti* populations exposed were resistant or showed suspected resistance to lambda (no information from Guatemala). Permethrin data, from El Salvador and Honduras, showed resistance of local populations to this insecticide (Figure 6).

**Figure 6.**
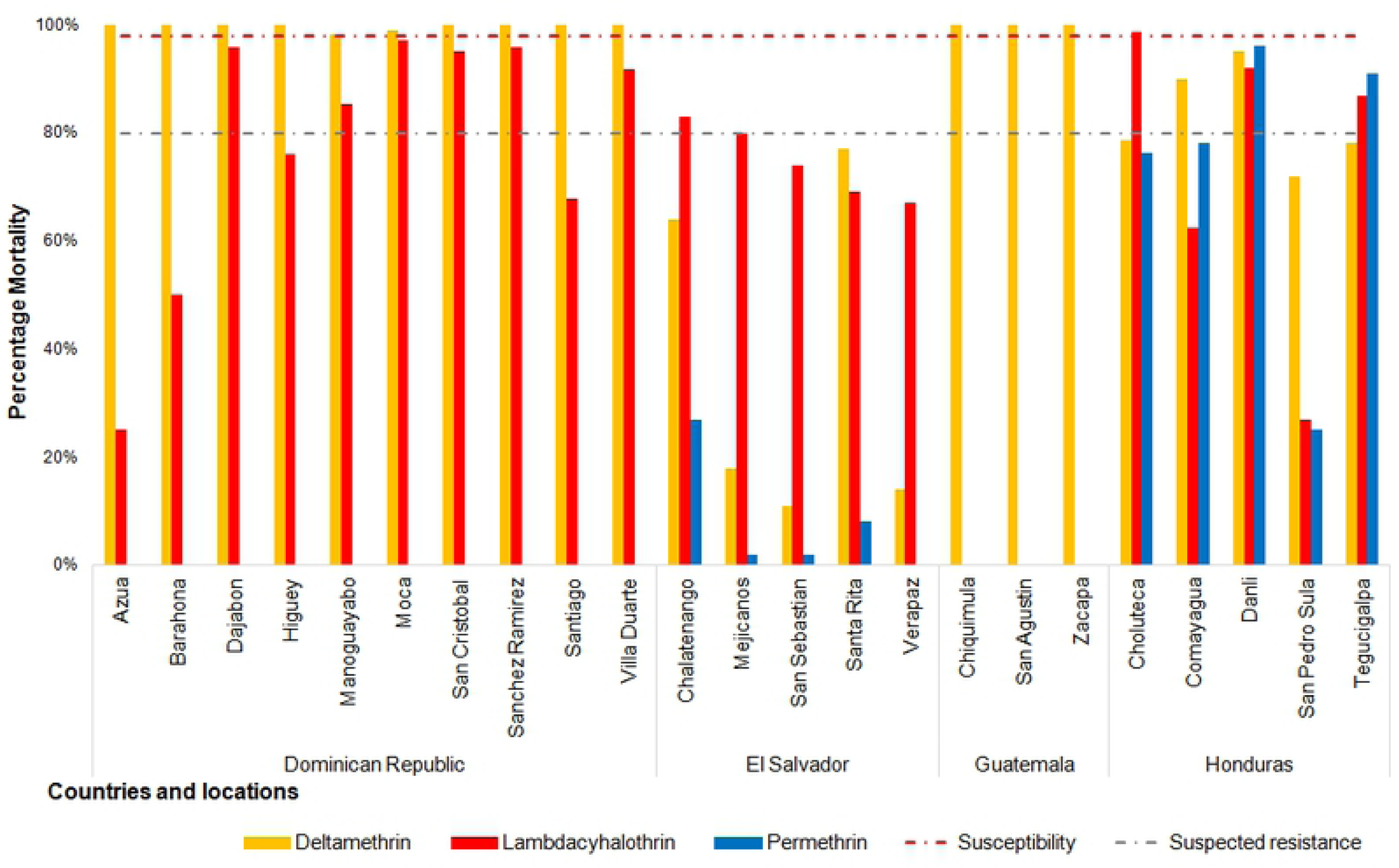
Mortality of *Aedes aegypti* populations from Dominican Republic, El Salvador, Guatemala and Honduras when exposed to five pyrethroids, using CDC diagnostic doses and 30 minutes diagnostic time. The horizontal red dotted line represents 98% mortality threshold which delimits susceptibility; results recorded between this threshold and the grey dotted line (80% mortality) are interpreted as suspected resistance, and results below the 80% mortality are indicative of resistance to the corresponding insecticide.

The organophosphate susceptibility tests demonstrated malathion resistance in only two Honduran sites (Comayagua and San Pedro Sula), with another two sites recorded as suspected resistance (Choluteca and Danli) in Honduras and Chalatenango in El Salvador. Remaining mosquito populations tested showed susceptibility to malathion (Figure 7). Wide spread resistance to pirimiphos-methyl was documented in El Salvador, with site specific variation in susceptible and suspected resistance in the Dominican Republic and Honduras (Figure 7).

**Figure 7.**
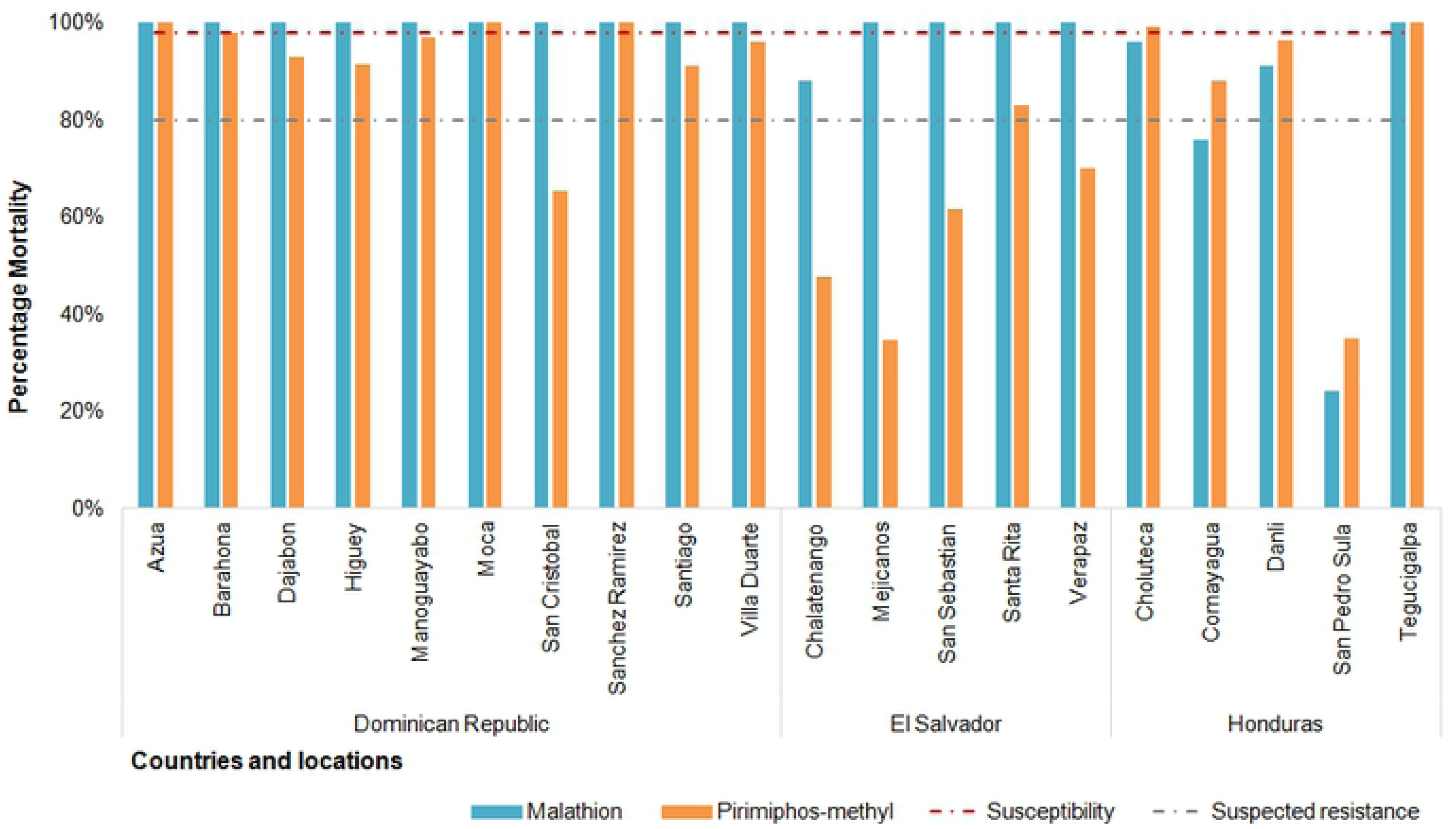
Mortality of *Aedes aegypti* populations from Dominican Republic, El Salvador and Honduras when exposed to organophosphates (malathion and pirimiphos-methyl), using CDC methodology. The horizontal red dotted line represents 98% mortality threshold which delimits susceptibility; results recorded between this threshold and the grey dotted line (80% mortality) are interpreted as suspected resistance, and results below the 80% mortality are indicative of resistance to the corresponding insecticide.

Finally, all mosquito populations across countries showed full susceptibility to the carbamate bendiocarb, except in the population from San Pedro Sula, from Honduras, which indicated suspected resistance (83% mortality).

### Comparison of WHO bioassays and CDC bottle bioassays

The comparison between the susceptibility status of *Ae. aegypti* populations as determined with the CDC and WHO methods varied according to the insecticide tested. The results for exposed mosquitoes from Honduras and El Salvador, consistently documented resistance to permethrin and deltamethrin with both methods (Tables 2 and 3). However, the status of susceptibility in sample populations from Guatemala and Dominican Republic, showed radical differences when exposed to deltamethrin. For the populations exposed to lambda-cyhalothrin, we recorded similarities in findings across all countries, with some populations found as susceptible (Table 3).

**Table 2.**
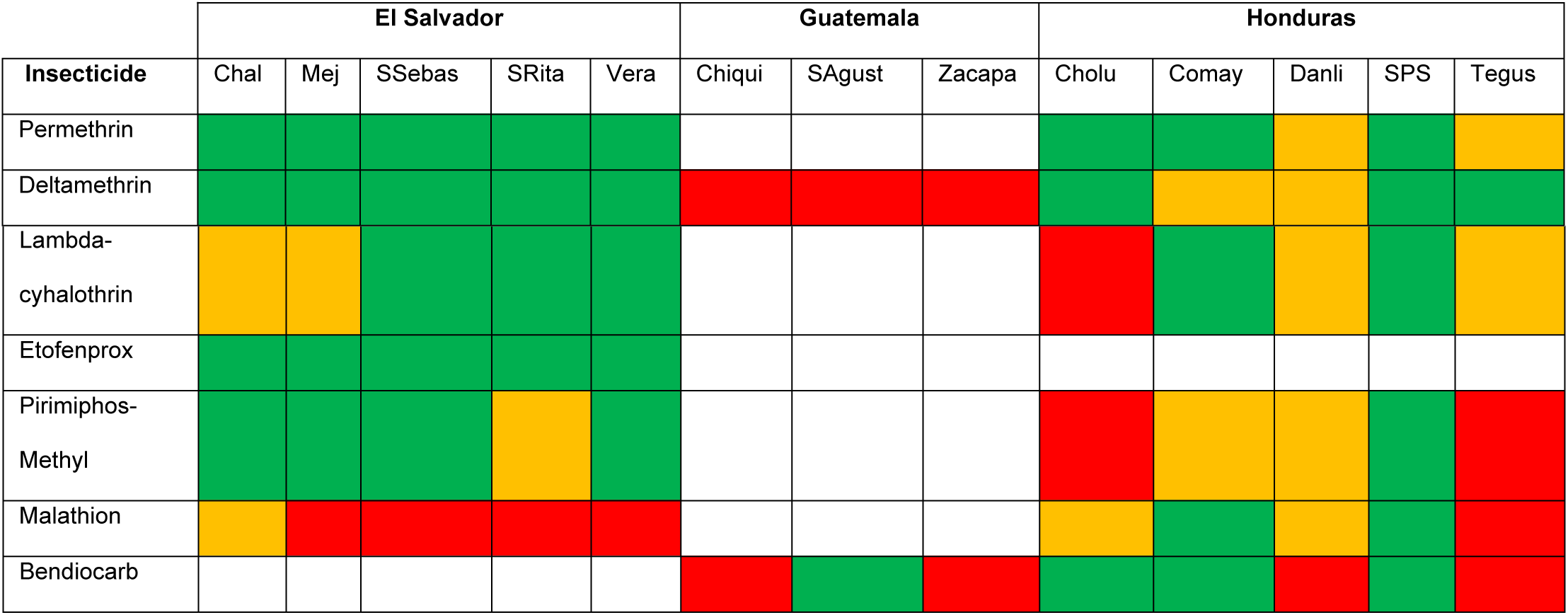
Comparison of susceptibility results in sample populations of Dominican Republic with the two standard methodologies to evaluate insecticide susceptibility (WHO vs. CDC). Green color= same; orange color= similar; red color= different; white color= tests were done with only one of the methodologies. Codes for study sites: Barah= Barahona; Dajab = Dajabon; Hig = Higuey; Mano = Manoguayabo; SCrist = San Cristobal; San Rami= Sanchez Ramirez; Santiago = Santiago de los Caballeros; VDuarte= Villa Duarte.

**Table 3.**
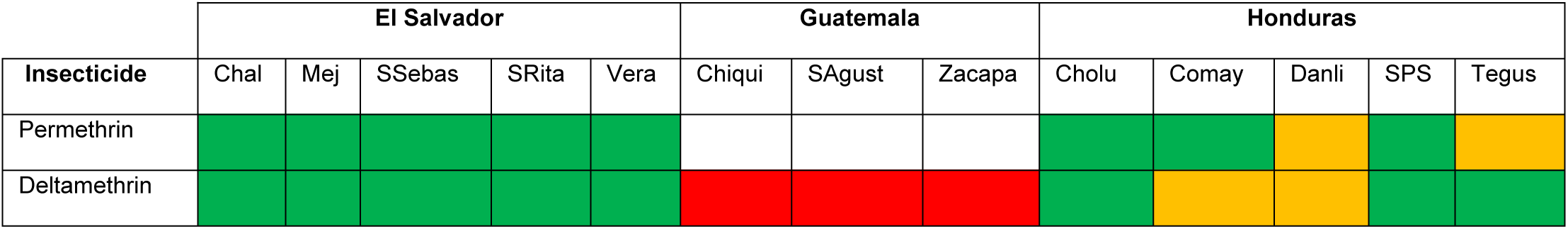
**Comparison of susceptibility results in sample populations of El Salvador, Guatemala and Honduras with the two standard methodologies to evaluate insecticide susceptibility (WHO vs. CDC)**. Green color= same; orange color= similar; red color= different; white color= tests were done with only one of the insecticides. Codes of site names: Chal= Chalatenango; Mej= Mejicanos; SSebas= San Sebastian; SRita= Santa Rita; Vera= Verapaz; Chiqui= Chuiquimula; SAgust= San Agustin; Cholu= Choluteca; Comay= Comayagua; SPS= San Pedro Sula; Tegus= Tegucigalpa. Note: In Guatemala only deltamethrin was used with both methods, while in Haiti no insecticide was used with both methods -alpha-cypermethrin was only carried out using the WHO method thus Haiti was not included in the comparison.

WHO and CDC assays testing the susceptibility of mosquito populations to the organophosphate malathion showed contrasting results for most of El Salvador and Dominican Republic samples and one site in Honduras. The recorded susceptibility of *Ae. aegypti* populations to Pirimiphos-methyl showed more congruent results among sites in El Salvador, with more contrasting sites in Honduras and the Dominican Republic. Finally, the comparison between the testing with the carbamate bendiocarb showed contrasting data for the Dominican Republic populations, while only two sites in Honduras and two sites in Guatemala showed incongruence in the data (Figure 8). In almost all the cases displayed in figure 8, where a difference or contrasting results were found with the two methodologies, CDC tests diagnosed susceptibility in the exposed populations while WHO tests diagnosed resistance or suspected resistance in the same site.

**Figure 8.**
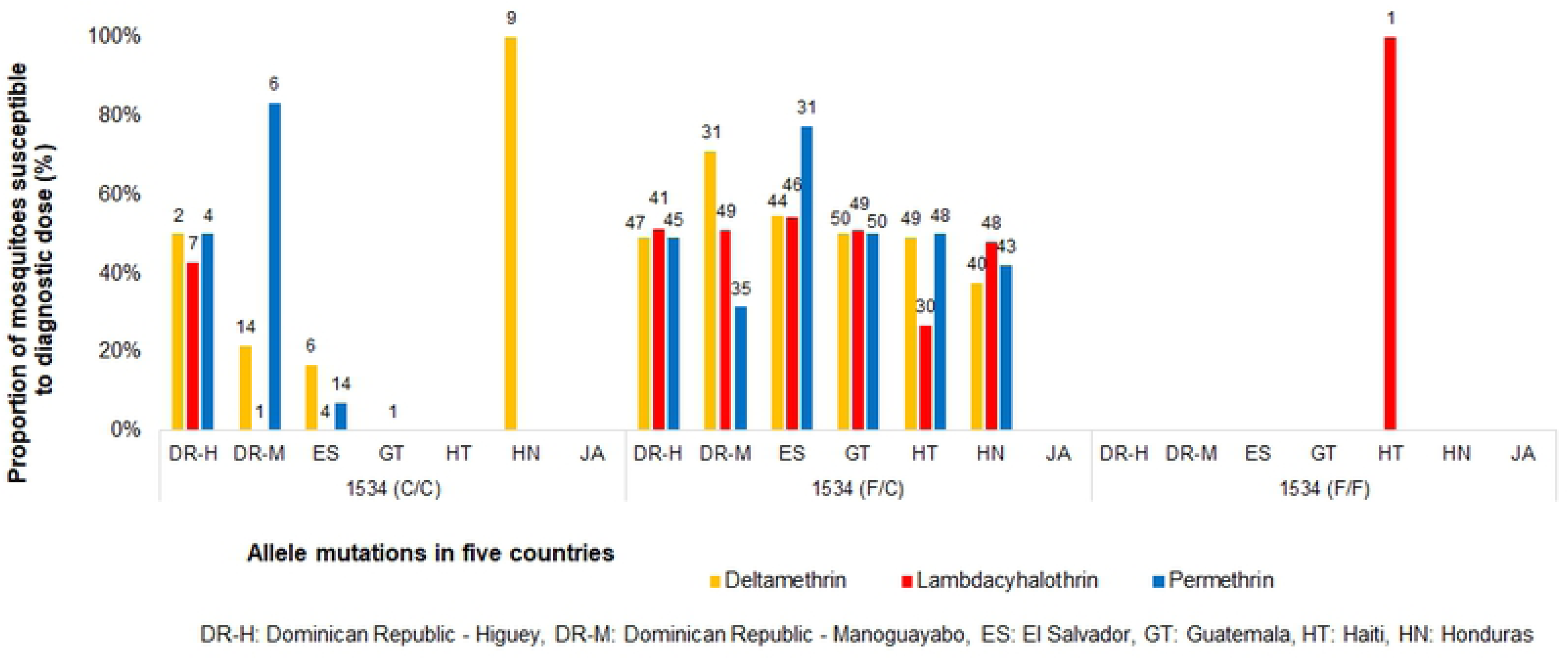
F1534C *kdr* genotyping in *Ae. aegypti* populations from countries of Central America and the Caribbean. The populations screened for the F1534C *kdr* mutation include Higuey (DR-H), Manoguayabo (DR-M), San Sebastian (ES), Chiquimula (GT), Dessources (HT) and Tegucigalpa (HD). The numbers on top of each bar are the number of mosquitoes that showed the respective genotype. In general, the presence of the V1016I mutated allele indicates a raise in tolerance to all three pyrethroids (Figure 9). All mosquitoes that were wild-type for V1016I were diagnosed as susceptible for lambda-cyhalothrin in both the Dominican Republic and the Haiti mosquito populations (Figure 9). Similarly, all wild-type mosquitoes in Honduras were susceptible to permethrin.

**Figure 9.**
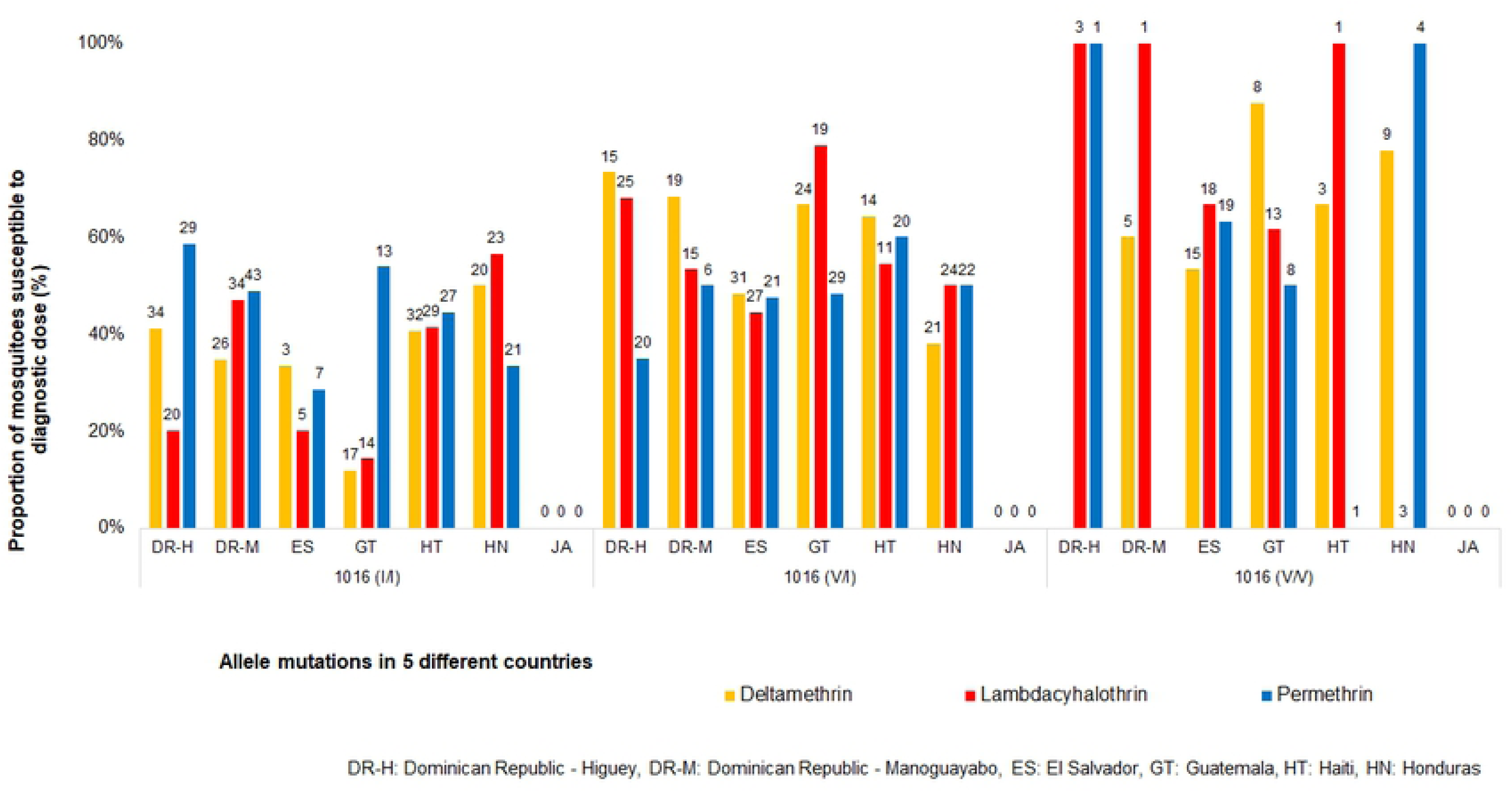
V1016I *kdr* mutation genotyping on *Ae. aegypti* population from countries of Central America and the Caribbean. The tested populations included Higuey (DR-H), Manoguayabo (DR-M), San Sebastian (ES), Chiquimula (GT), Dessources (HT) and Tegucigalpa (HD). The numbers on top of each bar are the number of mosquitoes that showed the respective genotype.

### *Kdr* genotyping

Target mutations studied comprised I1011V, I1011M, F1534C and V1016I. All samples (whether resistant or susceptible after WHO testing), were diagnosed as having the wild-type allele for I1011V. Forty-five of these samples, representing both survivors and non-survivors, were sequenced to ensure that the assay was functioning as expected. Sequence alignment are presented as supplementary information (S3). Sequencing determined that the processed samples had the wild-type allele for I1011V demonstrating the validity of the assay. These samples were also identified as wild-type for the I1011M allele - since they would have resulted in ‘wild-type’ results for the PCR assay (demonstrated definitively with the sequencing); therefore, these PCRs were not performed for I1011M and all mosquitoes were considered wild type for that allele too.

The majority of individuals across sampled countries were heterozygous for the F1534C mutation, regardless of its status of resistant or susceptible to any of the three insecticides (Figure 8). In Higuey (DR) there was a presence of homozygous mutation, but it was present in similar percentages in resistant and susceptible mosquitoes. A high number of homozygotes were found in Manoguayabo (DR) and Honduras, mainly related to susceptibility to permethrin and deltamethrin respectively. In contrast, all mosquitoes that were killed by lambda-cyhalothrin in Haiti were wild-type homozygous (Figure 8).

## Discussion

Despite dengue hyperendemicity in the Central America and Caribbean region, and the chikungunya and Zika epidemics, governments struggle to implement insecticide monitoring and surveillance programs to inform IVM activities. Several factors in the public health scenario of the sampled countries have resulted in gaps in mosquito surveillance, which translates into a lack of data on the insecticide susceptibility of *Ae. aegypti* mosquitoes towards decision-making. The complex scenario of public health involves competing demands for massive insecticide applications, due to political and social pressures, and the weak technical and logistical capacity of the national programs that deliver only sporadic and limited actions to prevent and control arboviruses transmission. Prior to the implementation of the Zika AIRS Project, none of the five countries included in this study had functioning programs that monitored insecticide resistance in Zika vectors. Additionally, this study follows the path of only few previous studies that have explored the molecular mechanisms of insecticide resistance present in the region (51–53) and is the only study that raises technical issues regarding the dual internationally-accepted system of detecting and reporting insecticide resistance that might result in contrasting outcomes.

### Insecticide resistance and intensity

Resistance to the pyrethroid permethrin is present in the vast majority of mosquito populations across countries, independently of the method of use. As a matter of fact, permethrin has been one of the most widely used insecticides to control *Ae. aegypti* due, in part, to the market availability and lower cost (54, 55). That is not the case with etofenprox, a pseudo-pyrethroid that is seldom employed in vector control operations in the region; however, resistance levels to etofenprox were also documented in this study. The selective pressure by permethrin (39) and possible cross-resistance mechanisms product of decades of use of DDT (56, 57) in the region are likely the cause of this widespread resistance. Additionally, permethrin is a type 1 pyrethroid, which can dissociate faster from the voltage-mediated sodium channels (VMSC), hence being more likely to originate resistance than pyrethroids type II, which block the channels several seconds longer (58).

The status of *Ae. aegypti* susceptibility to deltamethrin and lambda-cyhalothrin in the sampled countries was highly dependent of the method of testing. The CDC bottle assays demonstrated either susceptibility or suspected resistance, while WHO bioassays tended to result in resistance (contrary to previous research (59)).

Deltamethrin is a widely used insecticide in the whole region, applied mainly through ULV fogging and other types of spraying aiming to control adult populations of *Ae. aegypti*, so a resistant status was likely (11, 60, 61). Although lambda-cyhalothrin is not used as much as deltamethrin, examples of cross-resistance are reported in literature (62, 63), notwithstanding, both of them are type II alpha-cyano pyrethroids (64).

According to the results obtained with the WHO tests, resistance to the organophosphates malathion and pirimiphos-methyl is ubiquitous in the sampled *Ae. aegypti* populations. Malathion is considered the second choice of preference after deltamethrin, and is currently used in replacement of deltamethrin as a strategy for insecticide resistance management (60, 65, 66). Malathion has been also more recently been used in the Latin American region, so there is no surprise if loss of susceptibility is reflected in the results. Additionally, the region has been using the organophosphate temephos as the frontline chemical tool to control *Ae. aegypti* larvae, using several tons per year across countries (82 metric tons of organophosphates were used in larviciding between 2000-2009 in the Americas (67)). In a separate publication, parallel to this study, temephos resistance was also evaluated and is present in the Central American countries included here (data not included). Although this has previously been debated, it is possible that the selective pressure and resistance emergence in larvae can be transferred to adult mosquitoes against the same insecticide family (68–70). As pirimiphos-methyl is scarcely used in the region, resistance to this product could be a cross-resistance phenomenon stemming from the continuous and wide spread use of temephos (26, 71, 72). Resistance to malathion and pirimiphos methyl have been reported elsewhere (21, 73–75), and in the LAC region (26, 36, 71, 76–81).

Bendiocarb was the only carbamate tested in this study. Although it is not used in ULV applications, it may be used for indoor residual spraying (IRS) to deliver residual killing. IRS is being tested against *Ae. aegypti*, so the susceptibility level is particularly relevant. CDC bottle assays resulted in complete susceptibility in all mosquito populations, while WHO bioassays suggested some resistant populations in Honduras and widespread resistance in Dominican Republic (the only two countries to use bendiocarb with both methodologies). These contrasting results between CDC and WHO diagnostic doses resulting in confirmation of resistance or susceptibility being impossible, with the most conservative conclusion being that bendiocarb resistance is suspected.

WHO intensity bioassays performed in El Salvador and Honduras showed that *Ae. aegypti* populations tested in those countries have a high intensity resistance to permethrin, and moderate to high intensity resistance to deltamethrin and lambda-cyhalothrin. Resistance to pirimiphos methyl in both countries was classified as moderate. Monitoring the intensity of insecticide resistance regularly is essential in measuring goals of insecticide resistance management, to offer information on potential operational failure and optimize resources in a mosquito control program by selecting the most appropriate insecticides.

The documentation of susceptible mosquito populations to certain insecticides in the region, or that we might be overestimating deltamethrin resistance, and that CDC diagnostic dose is realistic, is hopeful. The susceptibility of different mosquito populations given a different biological and population genetics background is variable, therefore re-formulation of diagnostic doses obtained with different reference mosquito populations might be needed. Even more, a technique that links resistance with operational failure could be envisioned as essential insecticide product information for future formulations. In the face of uncertainty on the susceptibility status, the best choice is to follow an insecticide resistance management approach, continue routine monitoring and evaluating the insecticide products used in vector control operations (12, 82).

### Mechanisms of pyrethroid resistance: *kdr* screening

Mechanisms of insecticide resistance act in different ways: while target site mutations would result probably in knockdown and recovery (due to rapid dissociation of the insecticide molecules in the voltage-gated sodium channel), enzymatic resistance would probably result in mosquitoes tolerating the insecticide and not being knocked down. Since the WHO and the CDC methodologies main measures are mortality and knockdown respectively, exploring the presence of *kdr* mutations could contribute to explain the discordance between methods; however, in this study only mosquitoes used in WHO assays were genotyped, so the arguments definitely will need further validation.

All mosquitoes screened resulted as wild-type for the I1011V and I1011M mutations, and most of the mosquitoes screened were heterozygous for the F1534C mutation, regardless of the susceptible or resistant status to the WHO diagnostic doses of permethrin, deltamethrin and lambda-cyhalothrin. Interestingly, there was an increase of allelic frequency of the V1016I mutation in mosquitoes that survived the three pyrethroids across all countries. In *Ae. aegypti*, several *kdr* mutations have been linked with pyrethroid resistance. In particular the mutations D1763Y, F1534C, G923V, I1011M, I1011V, L982W, S989P, V1016G, V1016I, T1520I and V410L (29, 57, 83–86). In America, *kdr* mutations have been reported in *Ae. aegypti* populations from Ecuador, (22), United States (23, 24), Colombia (25, 26), México (27–29), Brazil (30–35), Lesser Antilles (36–40), French Guiana (37), Venezuela (29, 41), Cuba (29, 39), Panamá (42), Puerto Rico (43). This is the first report of *kdr* mutations for the countries included in this study, except for Haiti (87).

Resistance to pyrethroids in mosquitoes have been widely associated with the F1534C (21, 23, 25, 27, 29, 30, 32, 39–41, 83, 88–101) and V1016I mutations (24, 25, 27, 29, 30, 32, 36, 38, 40, 41, 88, 93, 97, 100, 102, 103). The simultaneous presence of both mutations has been associated with enhanced tolerance to deltamethrin in the past (31). However, the relationship between both mutations in relation to resistance is not clear. The fact that most of individuals are heterozygous for F1534C seems to indicate that its presence is not associated with resistance - although the theory that it is contributing to resistance in association with other mutations cannot be discarded. Other mutations such as G923V, (reported in the Americas (29, 57, 84, 86, 93)) and S989P, have been reported to be associated with resistance when in combination with other mutations (98, 99, 104), were not screened in this study but should be considered for future work.

### Differences between the WHO and CDC susceptibility classifications

When a country designs an insecticide resistance monitoring program, it usually selects one of the two available standardized methodologies: WHO or CDC. Based on those results, decisions are made on insecticide selection to guide public health program implementation. In the Latin American region, the CDC bottle assays are more commonly used mainly because the procurement process it’s easier. Conversely, WHO kits and insecticide impregnated papers are generally more difficult to obtain in the Americas because of geographic distance with Malaysia, language barriers, problematic procurement processes, etc. (59, 105). In addition, there are claims of quality loss of the impregnated papers in the transportation process. However, both methods are conventionally considered to be equally valid and hypothetically should offer similar information on mosquito susceptibility. In this study both methods were used on the same mosquito populations across 5 countries resulting in contrasting susceptibility classification. This is at best confusing and does not orient countries on which method to use. Thus it begs the question, is the information provided by each method different at its core, or does it refer to the same insecticide susceptibility concept?

WHO testing employs insecticide impregnated filter papers, diluted in oil (OPs and CA) or alcohols (PYR). The papers are impregnated with diagnostic doses, which are supposed to kill 100% of susceptible mosquitoes. Impregnated papers have an expiry date that lasts 1 year (106), and can be used 6 times maximum only. There are reports of loss of effectiveness of the insecticide impregnated papers only after 4 uses (107). Exposure time is likely to vary between each mosquito because there are areas of the WHO-kit cylinder that are not covered on insecticide (59); however, in the case of insecticides with repellent properties, mosquitoes tend to behave actively and fly within the cylinder, disturbing other mosquitoes and forcing exposure. CDC testing uses fully-coated insecticide bottles, so mosquitoes are continuously exposed no matter if they fly or not. All type of insecticides are diluted in alcohols and bottles are coated usually a few days or the same day before the test; according to the guidelines, organophosphates and carbamates degrade faster than pyrethroids. It is not clear how these methodological differences might affect the response of mosquitoes to the insecticide doses.

Comparing dosage equivalences in CDC bottle assays and WHO bioassays is close to impossible, because the insecticide is delivered in distinct ways (concentration on surface versus concentration percentage) and there is no way to measure how much insecticide an insect is actually exposed to. However, both methodologies are believed to offer the same basic dual outcome: resistance or susceptibility. This is where differences in the final outcome are problematic, even if it’s understood that there are methodological differences. Thus, the central problem is not that there are technical differences between both methods, but that the outcomes for the same mosquito population can be *different* (17).

One of the potential causes of difference resides in the original mosquito strains used for calibrating the diagnostic doses. As it was mentioned, each organization used several susceptible mosquito strains to test a range of insecticide concentrations and calculate the diagnostic dose (defined by WHO as the double of the lethal concentration 99, i.e. the double of the concentration that kills 99% of susceptible mosquitoes). Those mosquito populations (named Rockefeller, New Orleans, Liverpool, and others) had their own genetic and phenotypic background, and there are possibilities that they respond differently to the diagnostic concentrations than current natural mosquito populations. Also, some of those strains have been through several re-colonization processes, bottlenecks and inbreeding for decades (108, 109). Ideally, each country should establish a susceptibility baseline and monitor the evolution of resistance in comparison to that baseline, but the reality is that 1) there are virtually no mosquito populations that have not been exposed to insecticides or other type of xenobiotics and 2) it is likely that the capacity of governments to perform that task is not up to the task, at least for the immediate future.

There are other relevant questions that have been discussed in comparing both methodologies. For example, some differences such as the angle in which the WHO cylinders are kept during the bioassay might change the outcomes (15). The use of single diagnostic concentrations in a world where resistance to some insecticides is almost universal is barely informative. The employment of intensity diagnostic concentrations (5x and 10x) is a step forward, but it is likely that a deeper dive into understanding the nature of insecticide as a multi-dimensional biological treat will be needed in order to extrapolate knowledge into clear and practical actions to prevent and manage insecticide resistance.

The contrasting results obtained between WHO and CDC methodologies for IR testing in this study ask a vital and outstanding question: *What methodology and entomological endpoints can be standardized and are adequate to decide whether there is resistance to an insecticide towards decision-making?* Ideally, each country should have developed susceptibility baselines and calculated diagnostic concentrations based on those, but given the current distribution and level of resistance, and even the history of DDT usage, and potential cross-resistance, such a task is clearly challenging. Perhaps the best solution, is to use both methods, and if insecticide resistance is found for at least one of them, that result should be the conservative verdict. However, countries in the region barely have resources and capacity (funding, variability in testing procedures, required training, mosquito rearing facilities, bioassay degradation, etc.) to do just one of them This is a clear opportunity for which the regional and international health authorities should aim future studies and guidelines to support countries in the process of understanding the information coming from the available tools.

### Limitations

This study was performed over a period of time spanning 2.5 years. Insecticide resistance in mosquito populations is a highly plastic in nature – varying based on the insecticide selective pressure (frequency, type of insecticide, etc.), population genetics, and other factors, hence, it was expected that the presence and intensity of resistance demonstrated temporal variation.

As any multi-country study of this proportion, and despite the supervision and continuous training, it is possible that the quality of a small portion of the dataset did not meet stringent standards expected. However, since these bioassays were performed in optimal conditions and by trained people, and that they represent real data that goes towards decision-making for Ministries of Health, the outcomes and results are valid for all implementation partners. In addition, due to time constraints, human resources and insecticide priorities, all insecticides could not be tested using both methods in all sites across countries.

Only susceptible and resistant mosquitoes were genotyped in the WHO bioassays. Genotyping mosquitoes from the CDC bioassays could be a good opportunity to link target site mutations with the contrasting results between methodologies.

### Final recommendations

Based on these results, the Ministries of Health of El Salvador, Guatemala, Honduras, Haiti and Dominican Republic should establish national networks for insecticide resistance surveillance and management of *Ae. aegypti.* Additionally, a technical evaluation of the effectiveness of commercial insecticides for ULV deployment that contain any pyrethroid or organophosphate is urgently needed, particularly those containing permethrin or etofenprox and any insecticide that resulted in a mosquito population survival to the 10x diagnostic dose (high level of resistance). Ideally, these would include epidemiological indicators in addition to entomological ones typically used. The specific tools used to establish the insecticide resistance management networks, that can include insecticide rotations, mosaics or combinations with other molecules, must be discussed and standardized in an inter-disciplinary context with the technical support of international organizations such as the Pan American Health Association (PAHO), and according to the technical and logistical capacities of each country.

This study suggests the widespread nature of at least one mutation related to pyrethroid resistance in the region. Ministries of Health, in association with academic institutions and international collaboration should monitor V1016 I and F1534C mutation frequency on an annual basis. This would provide insight on the evolution of this mechanism of resistance through the years of an insecticide resistance management program. Other mutations reported elsewhere in the literature, and future sequencing studies with samples from the LAC region are needed to better understand the evolution, distribution and molecular determinants of resistance. A successful insecticide application program, by default, would change transmission and vector population dynamics – including IR. Monitoring and surveillance would enable the timely adaptation and implementation of appropriate methodologies or molecules that fight this evolving paradigm.

Organizations such as the CDC and WHO/PAHO should work collaboratively in the unified release of revised diagnostic doses and adjusted methodologies. The current doses for *Ae. aegypti* can result in contradicting results, which is at best confusing for the institutions making decisions in public health.

## Acknowledgements

We would like to acknowledge the generous support provided by Audrey Lenhart and Lucrecia Vizcaino from the Centers for Disease Control and Prevention, Insecticide Resistance and Vector Control Team, Center for Global Health, Division of Parasitic Diseases and Malaria, Entomology Branch, throughout the implementation period of the ZIKA AIRS project. Special thanks to all the Ministries of Health of El Salvador, Guatemala, Honduras, the Dominican Republic and Haiti, for their active collaboration and input. This study could not have been conducted without the administrative and finance team of the ZIKA AIRS project, and colleagues like Paula Wood, Kassim Mohammued, Alex Stanchfield, Carmen Vilanova, Carlos Cardenas, Richard Fisher and Patricio Murgueytio, who consistently supported procurement orders, distribution and customs release processes of all entomological tools and supplies for the various countries.

## Supporting Information captions

S1 Table. List of all locations, countries, dates, insecticides, methodologies and mortality values obtained.

S2 Table. Summary of type of insecticide and concentrations employed for bioassays.

S3 Text file. Laboratory protocols for the kdr mutation molecular screening.

S4 Table. Raw data containing the genotypes for each specimen processed during the molecular studies.

S5 Table. Summary of the mutations vs. codons found during the molecular studies.

